# Lipidome profiles of plasma microvesicles differ in experimental cerebral malaria, compared to malaria without neurological complications

**DOI:** 10.1101/2020.07.28.224170

**Authors:** Amani M Batarseh, Fatemeh Vafaee, Elham Hosseini-Beheshti, Alex Chen, Amy Cohen, Annette Juillard, Nicholas Henry Hunt, Michael Mariani, Todd Mitchell, Georges Emile Raymond Grau

## Abstract

Cerebral malaria (CM), a fatal complication of Plasmodium infection that affects children in sub-Saharan Africa and adults in South-East Asia, results from incompletely understood pathogenetic mechanisms, which include an excessive release of microvesicles (MV). Plasma MV levels have been found elevated in CM patients and in the experimental mouse model.

We compared lipid profiles in circulating MV purified from CBA mice infected with *P. berghei* ANKA (PbA), which causes CM, to those from *P. yoelii* (Py), which does not. Here we show that plasma MV produced at the time of CM differed dramatically from those from non-CM mice, in spite of identical levels of parasitaemia. Using high-resolution LCMS, we identified over 300 lipid species within 12 lipid classes. Total lysophosphatidylethanolamine (LPE) levels were significantly lower in PbA infection compared to uninfected mice, while they were unchanged in Py MV, and lysophosphatidylcholine (LPC) was more significantly reduced in PbA mice compared to the other two groups. These results suggest, for the time, that experimental CM is characterised by specific changes in lipid composition of circulating MV, pointing towards triglycerides (TG) especially docosahexaenoic acid (DHA 22:6) containing species, phosphatidylethanolamine (PE), LPC, LPE, and diacylglycerol (DG) as potential important players in CM pathogenesis.

## Introduction

Cerebral malaria (CM) is a major neurovascular pathology complicating *Plasmodium falciparum* infection, which still is a major public health issue worldwide. CM is characterised by unarousable coma, neurological deficits and neurological sequelae. This debilitating syndrome accounts for the majority of malaria-induced deaths annually^1-3^, (WHO 2019: https://www.who.int/news-room/fact-sheets/detail/malaria).

The pathogenetic mechanisms of CM are exceedingly complex and therefore incompletely understood. One approach to this problem is the use of mouse models of CM. Dynamic interactions between infected erythrocyte sequestration, host cell activation and inappropriate immuno-inflammatory responses have been extensively studied^4-9^. The murine CM model has limits^10^ but also numerous positive aspects^11- 16^, which makes it an invaluable tool^17^. A widely used model for CM is inbred CBA mice infected with *P. berghei* ANKA (PbA), which leads to fatal disease with cerebral pathology within 10 days^18^. Conversely, infection of CM-susceptible mice with *P. yoelii* (Py) leads to hyper-parasitaemia and anaemia but without neurological complications^19^. This syndrome following Py infection is referred to as non-cerebral malaria (NCM).

In addition to their established roles in cell-cell interactions, extracellular vesicles (EV) play an important role in CM pathogenesis^9,20,21^. Microvesicles (MV), previously called microparticles, are one of the 4 families of EV. Now recognised as major elements in cell-cell communications^22^, notably in the central nervous system^23^, they play essential roles in homeostasis and are active players in inflammatory and immunopathological conditions^24^, including CM^21^.

Recently, lipid subspecies/ families have emerged as important regulators of pathophysiological conditions *in vitro*^25^ and *in vivo*^26^. The roles of lipids are increasingly known in inflammation, immunoregulation, metabolism and cancer^27^, as well as in malaria parasite biology^28^. Their involvement in EV biology has been reviewed recently^29^.

The aims of this study were to determine whether MV produced during CM and NCM differed in terms of lipid composition, and to evaluate whether some lipid species could be correlated with pathogenesis and might be biomarkers of disease severity and/or targets for therapeutic intervention.

## Materials and Methods

### Mice and parasite inoculation

We confirm that all experiments were performed in accordance with relevant guidelines and regulations. All mice used in this study were handled according to protocols approved by the University of Sydney Animal Ethics Committee (approval numbers K20/7-2006/3/4434, 418 and 326). Female CBA mice, 7 weeks old, were purchased from the Animal Resources Centre (Canning Vale, Western Australia). Mice were fed a commercial rodent pellet diet and had access to water *ad libitum*. Experimental mice were studied under pathogen-free conditions and monitored daily. PbA was a personal gift from Prof Josef Bafort, Prinz Leopold Institute, Antwerpen, Belgium^30^ and Py a personal gift from Prof John Playfair, London^31^ to GEG. Parasite stabilates were prepared as previously described^32^ and stored in liquid nitrogen.

Three experimental groups of mice were studied: non-infected (n = 10), PbA-infected (n = 7) and Py-infected (n = 8). Infection was induced by intra peritoneal injection of 1 × 10^6^ infected erythrocytes^32,33^. Mice were euthanized seven days post inoculation. Parasitaemia was monitored by counting 500 erythrocytes in Diff-Quick-stained thin blood smears.

### Blood sampling and MV preparation

Mouse venous blood was collected by retro-orbital venepuncture under anaesthesia into 0.129 mol/L sodium citrate (ratio of blood to anticoagulant 4:1). Samples were centrifuged at 1 500 g for 15 min at room temperature. Harvested supernatant was further centrifuged at 18 000 g for 4 min, twice, to achieve platelet-free plasma (PFP) and MV pellets. MV numbers were assessed as previously described^9^.

### Lipid extraction

Lipids were extracted from 1 mL of PFP following the MTBE protocol of Matyash *et al*.^34^. In brief, 300 μL of methanol containing 5 μL Splash Lipidomix deuterated standard (Avanti, USA – purchased from Sigma, Australia) was added to the MV pellets in Eppendorf tubes cooled on ice. Samples were vortexed briefly and incubated on ice for 10 min. MTBE (1 000 μL) was added to the tubes, which then were vortexed and the contents allowed to mix on a rotating shaker at 4 °C for 1 hour. Optima level H_2_O (250 μL) was added before samples were vortexed briefly and kept on ice for 10 min to allow phase separation. Following this, samples were centrifuged for 10 min at 10 000 g in a tabletop centrifuge set to 4 °C and 900 μL of the MTBE/Methanol top phase was transferred to 1.5 mL Eppendorf tubes. Lipid extracts were stored at −80 °C until analysed by liquid chromatography-mass spectrometry (LCMS).

### Liquid-chromatography mass spectrometry

For liquid chromatography, 900 μL aliquots of the lipid extracts were dried in a speed vacuum and reconstituted in 100 μL of isopropanol:methanol (2/1 v/v), vortexed for 20 sec twice and centrifuged at 10 000 g for 30 sec. Lipid extracts were transferred to glass vials with glass inserts and Teflon caps prior to analysis. Reversed-phase, ultra-high performance liquid chromatography (RP-UHPLC) was performed using a Vanquish liquid chromatography (LC) system (Thermo Fisher Scientific, Scoresby, VIC, Australia) fitted with a C30 column (Acclaim 2.1 × 150 mm, 3 μm particle size, Thermo Fisher Scientific, Scoresby, VIC, Australia) held at 10 °C. Two mobile phases were used; A: acetonitrile/water (60/40 v/v), 10 mM ammonium formate + 0.1% (v/v) FA and B: isopropanol: acetonitrile (90/10 v/v), 10mM ammonium formate + 0.1% (v/v) FA. For LC-MS operation, 5 μL of sample was injected onto the column with a solvent flow rate of 400 μL min^-1^. Mass spectrometry (MS) was performed on a Fusion Orbitrap mass spectrometer using targeted and untargeted lipidomics approaches (Thermo Fisher Scientific, Scoresby, VIC, Australia). LipidSearch software was used to annotate and quantify lipid species.

### Nomenclature

The lipid nomenclature used here is guided by literature recommendations of Liebisch *et al*.^35^ except for cholesteryl ester, which is abbreviated as ChE.

### Statistical analysis

Preprocessing and differential analyses were performed in R using the ‘*limma’* package. Raw lipid profiles were log2 transformed and normalised to equalise median absolute values across samples (see Supplementary Fig. S1 for *pre-* vs *post-*normalisation profiles). Moderated *t-*test^36^ was applied to normalised profiles to rank lipid species in order of evidence for differential expression; p-values were adjusted for multiple hypothesis testing using false discovery rate (FDR) correction. For correlation analysis, invariant lipids—i.e., those with interquartile range, IQR ≤ 1— were removed (Supplementary Fig. S2); 121 lipid ions out of 302 were retained for subsequent analysis. Pair-wise Pearson correlation was performed and the correlation matrix was visualised using the ‘*corrplot’* R package, where lipids were ordered using hierarchical clustering with ‘*complete’* agglomeration method.

## Results

### Qualitative and quantitative changes in MV produced in CM versus NCM

MV were purified and lipids extracted, as described, from the three groups of control, PbA-infected (i.e., with CM) and Py-infected (i.e., NCM) mice. Compared to those from controls, MV from PbA-infected mice showed a doubling of their proportion of of triglycerides (TG), a 25% reduction (47.6 *versus* 62.6%) in their cholesteryl ester (ChE) proportion, and a 50% reduction (2.8 *versus* 5.6%) in their lysophosphatidylcholine (LPC) content (Fig. 1A). These MV also presented a threefold increase (2.2 *versus* 0.7%) in their diacylglycerol (DG) content. Conversely, MV from Py-infected mice did not show such differences when compared to those from uninfected control mice. Parasitaemia levels were not significantly different between PbA- and Py-infected animals (not shown)^33^.

**Fig. 1.**
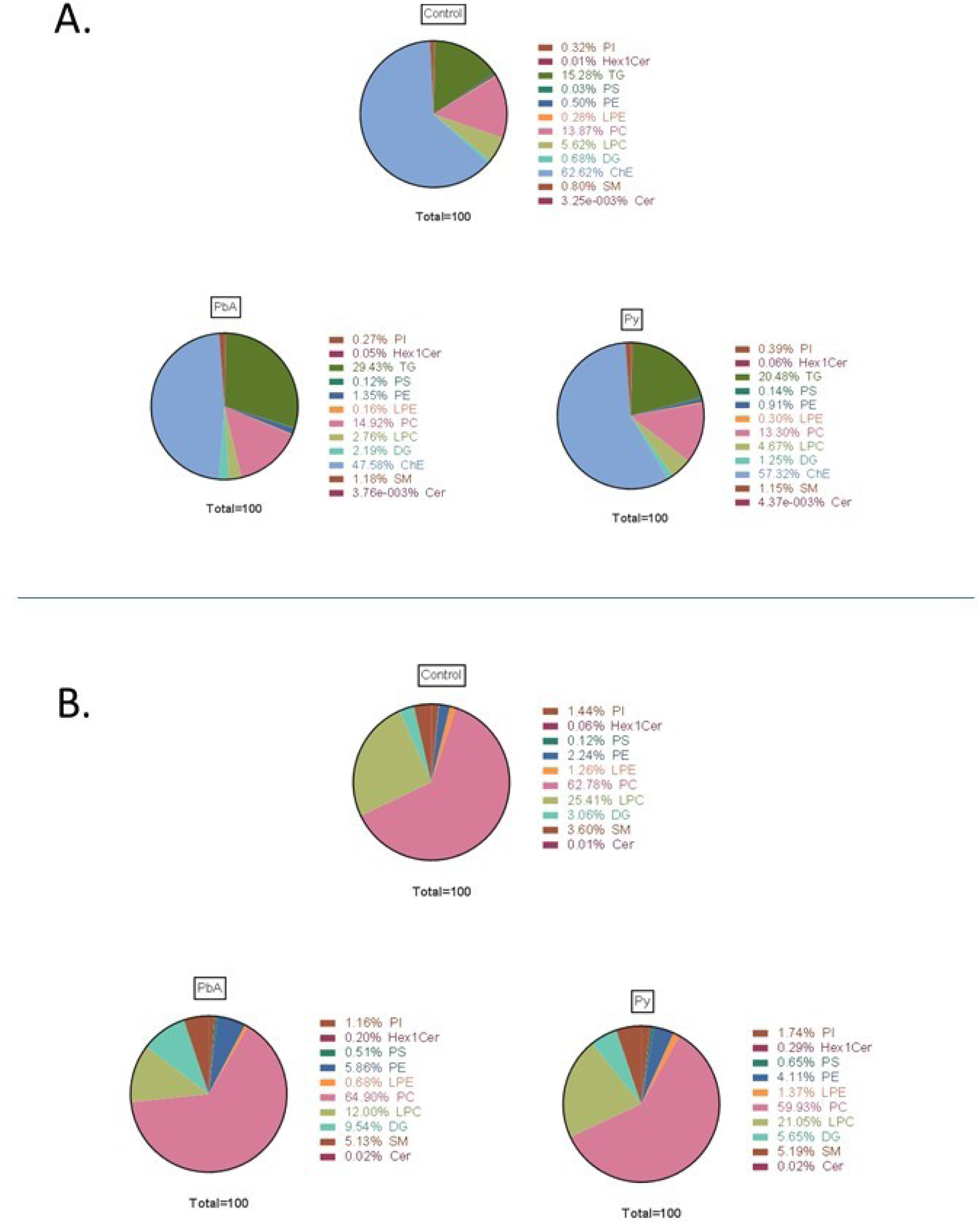
Lipid classes in plasma microvesicles (MV) from uninfected controls, PbA-infected and Py-infected mice. (A) All lipid classes. (B) levels without triglycerides (TG) and cholesteryl esters (ChE).

There have been reports suggesting that cholesteryl ester and TG are contaminants in MV preparations^25^. The levels of TG and ChE in the preparations are higher than other lipid classes, therefore we also analysed the results without these two classes of lipid (Fig. 1B). Under these criteria, the reduction in LPC in MV from PbA-infected animals was confirmed and a 3-fold increase in Hex1Cer was disclosed as well as a 5-fold increase for Py-infected mice.

### CM caused significant alterations in MV-derived lipid class composition

Significant changes were observed in the levels of DG, LPE, PE, LPC, PS and PI lipid classes in MV from PbA-infected vs control mice (Fig. 2). A quantitative analysis of these differences identified that, of these classes, LPC and LPE were decreased. In MV from Py-infected vs control mice, and DG, Hex1Cer, PE, LPC and PS lipid class levels were modified, while only LPC decreased. When comparing MV from PbA-infected to those from Py-infected mice, only DG, Hex1Cer, LPC and LPE amounts were significantly different. Interestingly, the amounts of both LPC and LPE lipid classes were more markedly reduced in MV from PbA-infected mice.

**Fig. 2.**
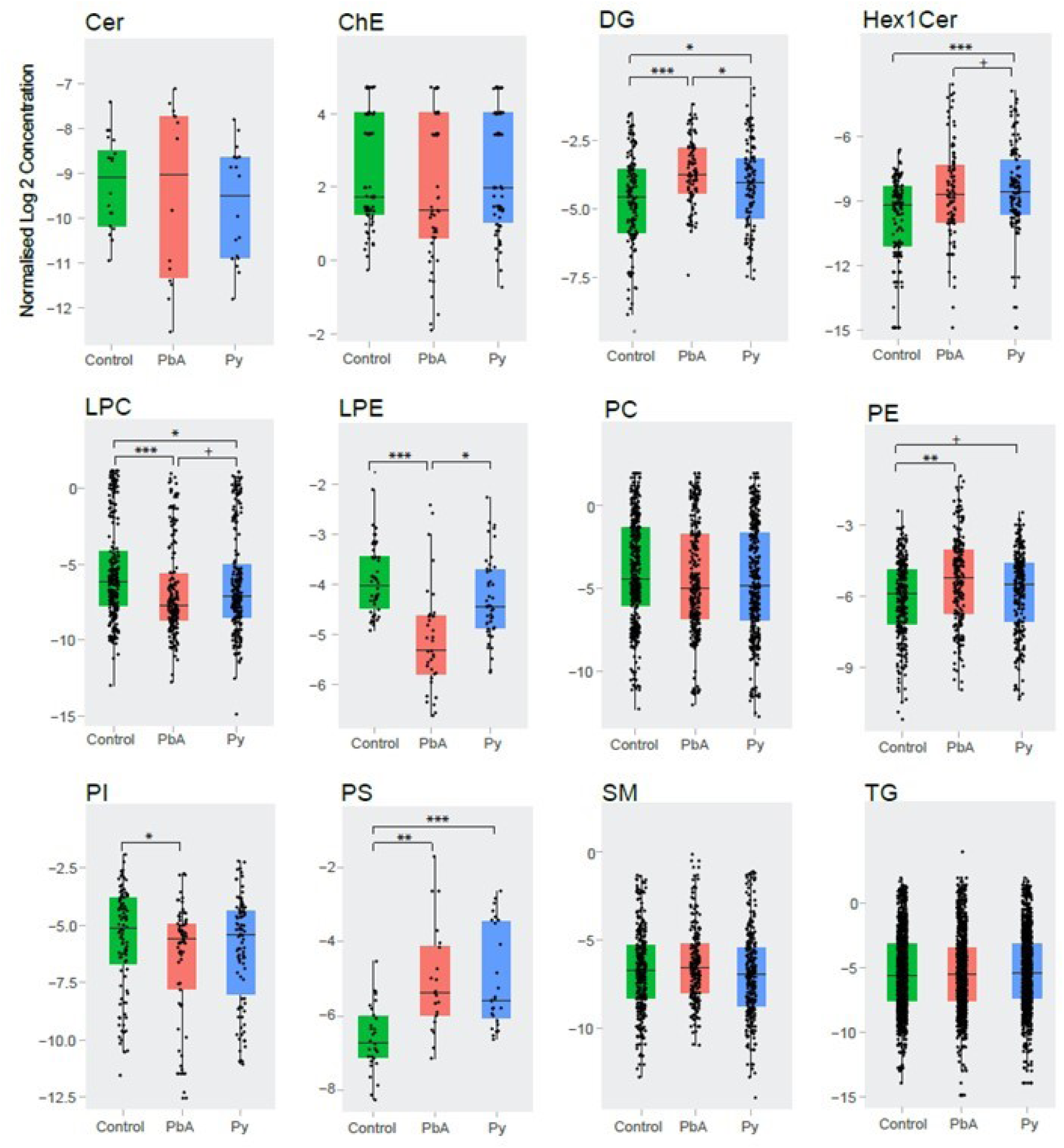
Altered lipid class levels in plasma MV from the three groups of mice. Boxplots showing comparisons of total lipid classes concentrations (pmol/ ul) between the groups of Control (green), PbA (red) and Py (blue). Data were log transformed and significance calculated using Students’ *t*-test. Significance annotation: p<0.05 +, p<10e-2 *, p<10e-3 **, p<10e-4 ***. Abbreviations as in Fig 1.

A principal component analysis (PCA) of MV lipidomes in the three groups of mice illustrates that MV in PbA were the most different from MV in control conditions (Fig. 3A). The numbers of *differentially expressed* lipids (i.e., adjusted p-value < 0.01 and |log2 fold-change|>1) in MV from the three categories of mice were further visualised using volcano plots and Venn Diagrams (Fig. 3B-D). MV from the PbA group differed from MVs from the Py group, with both increased and decreased lipid species (Fig. 3B and 3C). Volcano plots demonstrated that the lipidome of PbA-infected mouse MVs was dramatically different compared to controls, while that of Py-infected mice was not (Fig. 3B). When compared to controls, the abundance of 29 lipid species was increased in PbA-infected mice MV and 52 lipid species were decreased. In contrast, MVs from Py-infected mice showed only 9 lipid species increased and 2 lipid species decreased. Interestingly, when compared to those from Py-infected MVs, PbA had 20 increased lipid species and 30 decreased lipid species (Fig. 3C). The Venn diagram demonstrates that the lipidome of PbA-infected MVs was the most strikingly modulated (Fig. 3D). It also shows substantial overlap among differentially modulated lipids in ‘PbA vs Py-infected animals’ and ‘PbA-infected vs uninfected control mice’, which suggests similarity of the lipidomic profiles in Py-infected and uninfected mice when compared with PbA.

**Fig. 3.**
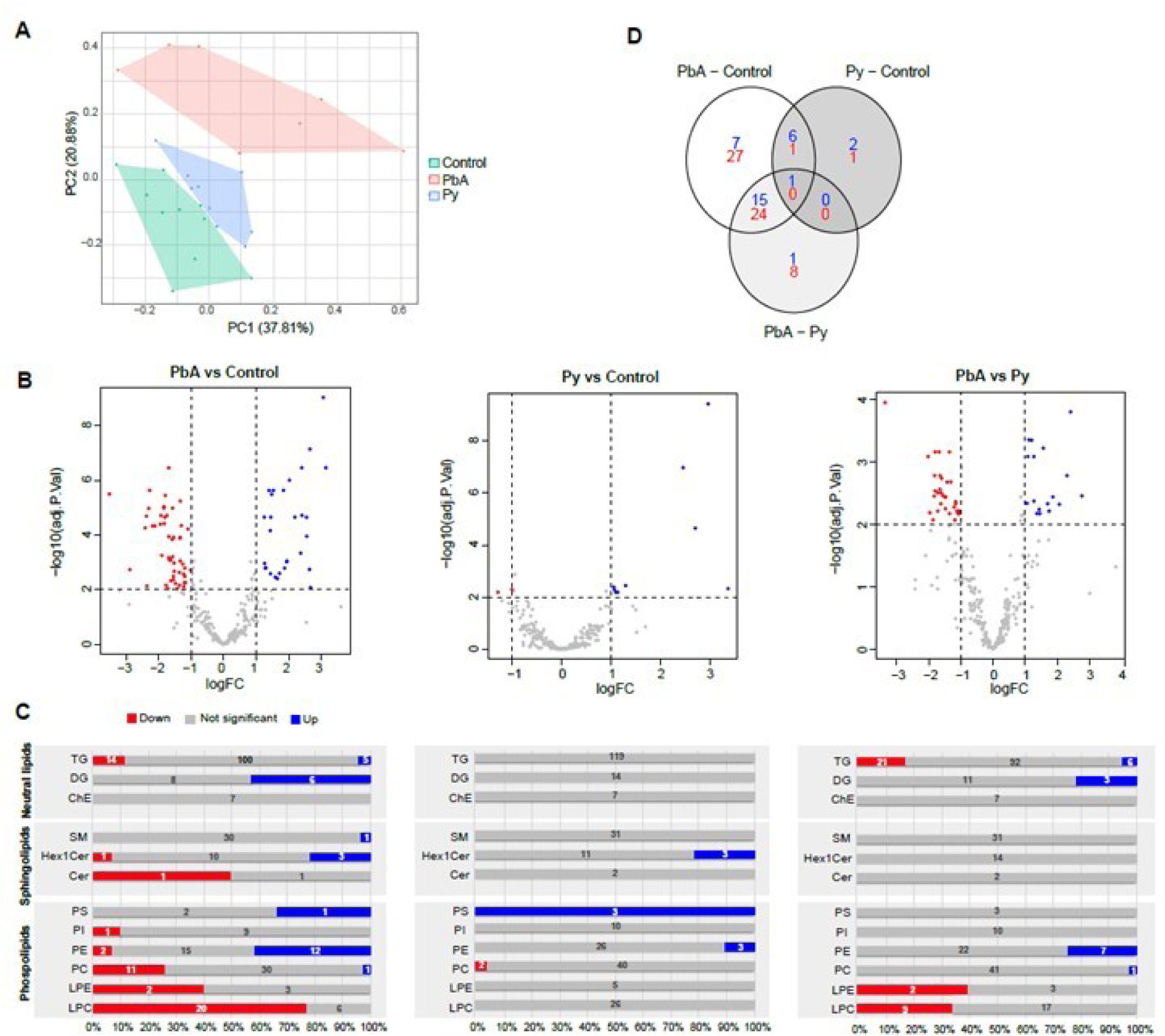
Differentially expressed lipids among MVs from the three categories. (A) Principal component analysis (PCA) of identified lipids (pmol/μL). (B) Volcano plots showing differentially expressed lipids in the various pairwise comparisons. (C). Bar charts representing the proportion of differentially expressed (DE) ions across each lipid graph. Blue, Grey and Red bars show the percentage of elevated, not significant, and decreased level of lipid molecules in each type. Numbers on top show the actual number of DE ions in each type. (D) Venn diagram of the distribution of differentially expressed lipids in the various comparisons. Significance is based on two-fold increase or decrease in lipid amounts plus an adjusted p-value <0.01 based on moderated t-test and fale discovery rate (FDR) correction.

### Quantitative analysis of lipid species among measured lipid classes

Comprehensive analysis of the lipid species was performed within each detected lipid class of the isolated MV-derived lipid extracts, and the composition of the lipid species was characterised. Tables I-III show the differentially expressed lipid ions in MV from three groups of comparisons, PbA-infected vs control mice (Table I), Py-infected vs control mice (Table II), and PbA- vs Py-infected mice (Table III). Lipid species with a positive fold change are shown in blue, and negative fold change in red.

**Table I.** Individual lipid molecules found at significantly different levels in PbA MV *versus* control. Ions elevated (decreased) in PbA are coloured in blue (red). Log2 fold-change and adjusted p-values are listed (i.e., moderated t-test and FDR correction). Ions are grouped into neutral lipids, sphingolipids and phospholipids and sorted alphabetically within each group.

| Lipid Ion | Log FC | Adjusted p-value | Lipid Ion | Log FC | Adjusted p-value | Lipid Ion | Log FC | Adjusted p-value |
| --- | --- | --- | --- | --- | --- | --- | --- | --- |
| DG(16:0_22:6) | 3.15 | 3.45E-07 | Hex1Cer(d18:1_16:0) | 3.06 | 9.36E-10 | PC(16:0_20:5) | -1.01 | 0.00189 |
| DG(18:1_20:4) | 2.19 | 2.22E-05 | Hex1Cer(d41:1) | -2.36 | 0.007303 | PC(O-16:0_20:4) | -1.34 | 0.00013 |
| DG(18:1_20:5) | 2.37 | 0.000461 | Hex1Cer(d42:1) | 2.67 | 0.00836 | PC(O-16:0_22:4) | -1.23 | 0.002657 |
| DG(18:1_22:6) | 1.72 | 0.002531 |  |  |  | PC(16:1_20:4) | -1.3 | 4.54E-05 |
| DG(18:2_20:4) | 2.41 | 1.90E-05 | LPC(14:0) | -1.69 | 0.000114 | PC(17:0_18:2) | -1.1 | 6.07E-05 |
| DG(18:2_22:6) | 1.95 | 0.000892 | LPC(15:0) | -1.83 | 3.83E-05 | PC(18:0_22:6) | 1.28 | 0.001628 |
| TG(16:0_16:0_22:6) | 1.94 | 0.000949 | LPC(O-16:0) | -1.19 | 0.003254 | PC(O-18:1_20:4) | -1.57 | 1.04E-05 |
| TG(16:0_18:1_23:0) | -1.19 | 0.001628 | LPC(O-16:1) | -1.54 | 0.004291 | PC(18:2_18:2) | -1.95 | 3.99E-05 |
| TG(16:0_18:1_24:0) | -1.54 | 0.003227 | LPC(16:1) | -1.84 | 1.04E-05 | PC(19:0_18:2) | -1.69 | 3.45E-07 |
| TG(16:0_20:4_22:6) | 2.56 | 0.000112 | LPC(17:0) | -1.34 | 0.000123 | PC(20:4_22:6) | -1.17 | 0.0055 |
| TG(16:0_22:6_22:6) | 2.64 | 0.001803 | LPC(O-18:1) | -1.68 | 0.000668 | PC(22:0_18:2) | -1.57 | 0.000624 |
| TG(O-16:0_16:0_18:2) | -1.2 | 0.005488 | LPC(18:1) | -1.58 | 0.000144 | PE(16:0_18:1) | 2.03 | 9.88E-07 |
| TG(O-16:0_18:1_18:2) | -1.36 | 0.006879 | LPC(18:2) | -2.11 | 4.64E-05 | PE(16:0_18:2) | 1.4 | 2.35E-06 |
| TG(18:0_18:1_22:0) | -1.62 | 0.004668 | LPC(18:4) | -2.28 | 2.35E-06 | PE(16:0_20:4) | 1.53 | 2.35E-06 |
| TG(18:1_20:4_22:6) | 1.88 | 0.001628 | LPC(19:0) | -1.75 | 1.94E-05 | PE(16:0_20:5) | 1.44 | 0.002587 |
| TG(18:1_22:0_18:1) | -1.71 | 0.009094 | LPC(20:0) | -1.81 | 2.24E-05 | PE(16:0_22:6) | 1.84 | 2.33E-06 |
| TG(18:1_22:0_22:0) | -1.9 | 0.000559 | LPC(20:1) | -1.84 | 9.10E-06 | PE(O-16:1_18:2) | -1.42 | 0.009094 |
| TG(18:2_20:4_22:6) | 1.59 | 0.003492 | LPC(20:2) | -2.3 | 1.06E-05 | PE(O-16:1_22:4) | 1.26 | 0.001092 |
| TG(24:1_18:2_18:1) | -1.77 | 0.006916 | LPC(20:4) | -1.54 | 0.000123 | PE(18:0_20:4) | 1.45 | 2.22E-05 |
| TG(24:1_18:2_18:2) | -1.48 | 0.005911 | LPC(20:5) | -2.37 | 2.04E-05 | PE(18:0_20:5) | 1.44 | 6.91E-05 |
| TG(25:0_18:1_16:0) | -1.26 | 0.001152 | LPC(22:1) | -1.23 | 0.007528 | PE(18:0_22:6) | 2.66 | 7.25E-08 |
| TG(25:0_18:1_18:1) | -1.63 | 0.000827 | LPC(22:4) | -2.41 | 5.44E-05 | PE(O-18:1_18:2) | -1.36 | 0.002351 |
| TG(26:0_18:1_16:0) | -1.52 | 0.001061 | LPC(22:5) | -2.18 | 4.64E-05 | PE(18:1_18:1) | 1.3 | 0.001572 |
| TG(26:0_18:1_18:1) | -1.67 | 0.000815 | LPC(22:6) | -1.33 | 0.000892 | PE(18:1_20:4) | 1.25 | 2.22E-05 |
| TG(30:1_18:2_20:1) | -3.51 | 3.19E-06 | LPE(18:1) | -1.79 | 3.58E-06 | PE(18:1_22:6) | 1.48 | 3.26E-06 |
| SM(d36:0) | 1.65 | 0.003903 | LPE(18:2) | -1.92 | 1.94E-05 | PI(O-34:3) | -2.89 | 0.001838 |
|  |  |  | PC(15:0_18:2) | -1.33 | 5.73E-06 | PS(18:0_22:6) | 2.4 | 3.45E-07 |

**Table II.**
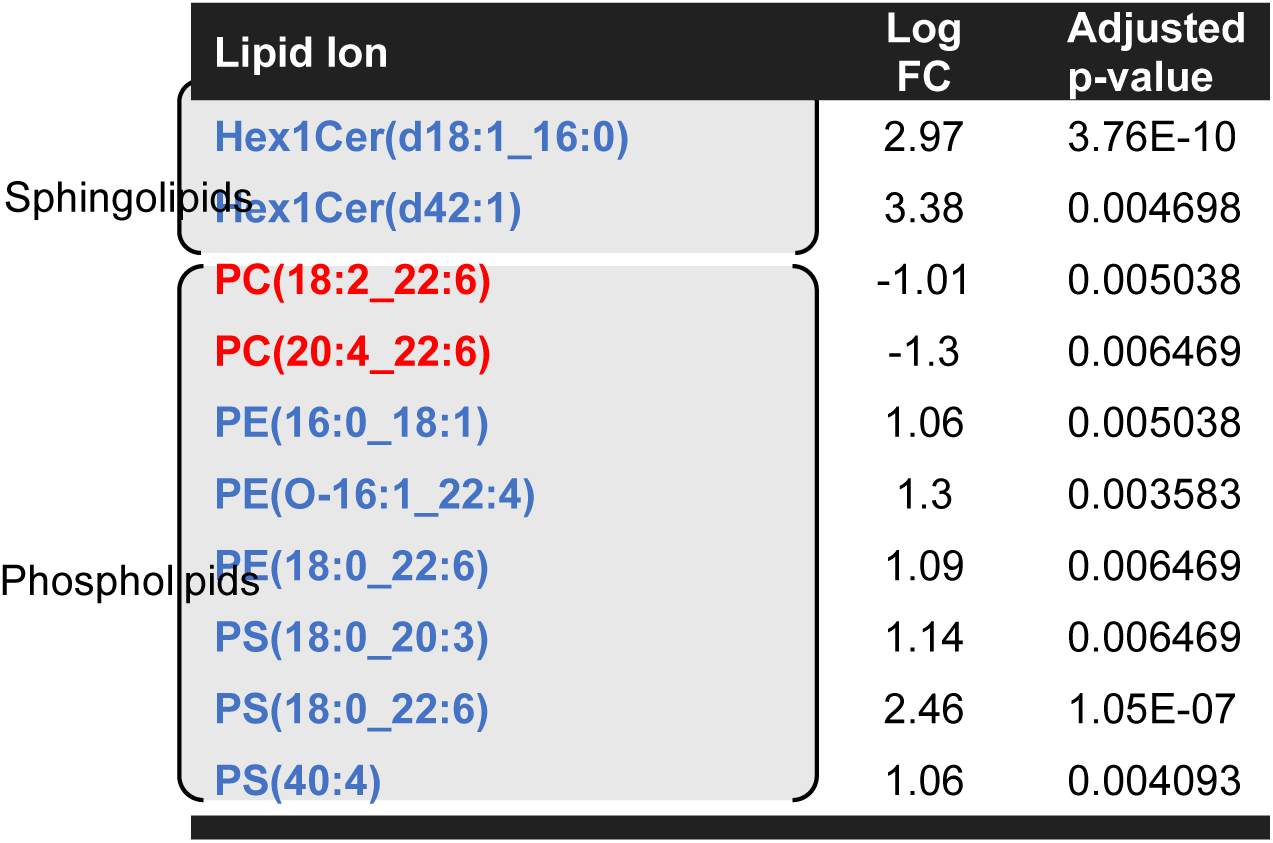
Individual lipid molecules found at significantly different levels in Py MV *versus* control. Ions elevated (dropped) in Py are coloured in blue (red). Log2 fold-change and adjusted p-values are listed (i.e., moderated t-test and FDR correction). Ions are grouped into sphingolipids and phospholipids and sorted alphabetically within each group.

|  | Lipid Ion | Log FC | Adjusted p-value |
| --- | --- | --- | --- |
| Sphingolipids | Hex1Cer(d18:1_16:0) | 2.97 | 3.76E-10 |
|  | Hex1Cer(d42:1) | 3.38 | 0.004698 |
| Phospholipids | PC(18:2_22:6) | -1.01 | 0.005038 |
|  | PC(20:4_22:6) | -1.3 | 0.006469 |
|  | PE(16:0_18:1) | 1.06 | 0.005038 |
|  | PE(O-16:1_22:4) | 1.3 | 0.003583 |
|  | PE(18:0_22:6) | 1.09 | 0.006469 |
|  | PS(18:0_20:3) | 1.14 | 0.006469 |
|  | PS(18:0_22:6) | 2.46 | 1.05E-07 |
|  | PS(40:4) | 1.06 | 0.004093 |

**Table III.**
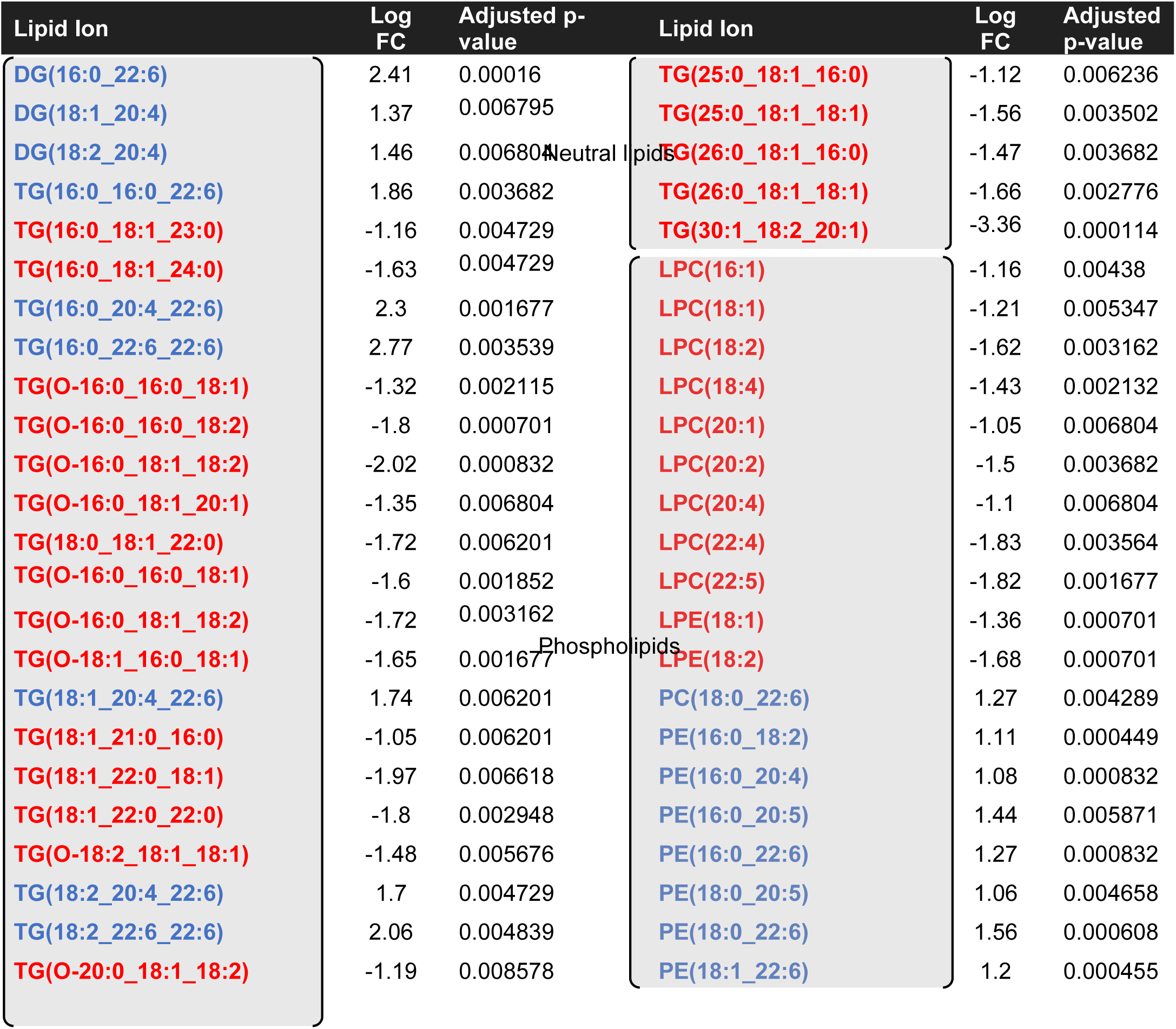
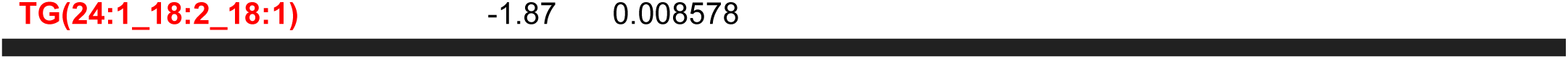
Individual lipid molecules found at significantly different levels in PbA MV *versus* Py. Ions elevated (dropped) in PbA are coloured in blue (red). Log2 fold-change and adjusted p-values are listed (i.e., moderated *t-*test and FDR correction). Ions are grouped into *neutral lipids* and *phospholipids* (located in the tables in a vertical fashion near each group of lipids) and sorted alphabetically within each group.

| Lipid Ion | Log FC | Adjusted p-value | Lipid Ion | Log FC | Adjusted p-value |
| --- | --- | --- | --- | --- | --- |
| DG(16:0_22:6) | 2.41 | 0.00016 | TG(25:0_18:1_16:0) | -1.12 | 0.006236 |
| DG(18:1_20:4) | 1.37 | 0.006795 | TG(25:0_18:1_18:1) | -1.56 | 0.003502 |
| DG(18:2_20:4) | 1.46 | 0.006804 | TG(26:0_18:1_16:0) | -1.47 | 0.003682 |
| TG(16:0_16:0_22:6) | 1.86 | 0.003682 | TG(26:0_18:1_18:1) | -1.66 | 0.002776 |
| TG(16:0_18:1_23:0) | -1.16 | 0.004729 | TG(30:1_18:2_20:1) | -3.36 | 0.000114 |
| TG(16:0_18:1_24:0) | -1.63 | 0.004729 | LPC(16:1) | -1.16 | 0.00438 |
| TG(16:0_20:4_22:6) | 2.3 | 0.001677 | LPC(18:1) | -1.21 | 0.005347 |
| TG(16:0_22:6_22:6) | 2.77 | 0.003539 | LPC(18:2) | -1.62 | 0.003162 |
| TG(O-16:0_16:0_18:1) | -1.32 | 0.002115 | LPC(18:4) | -1.43 | 0.002132 |
| TG(O-16:0_16:0_18:2) | -1.8 | 0.000701 | LPC(20:1) | -1.05 | 0.006804 |
| TG(O-16:0_18:1_18:2) | -2.02 | 0.000832 | LPC(20:2) | -1.5 | 0.003682 |
| TG(O-16:0_18:1_20:1) | -1.35 | 0.006804 | LPC(20:4) | -1.1 | 0.006804 |
| TG(18:0_18:1_22:0) | -1.72 | 0.006201 | LPC(22:4) | -1.83 | 0.003564 |
| TG(O-16:0_16:0_18:1) | -1.6 | 0.001852 | LPC(22:5) | -1.82 | 0.001677 |
| TG(O-16:0_18:1_18:2) | -1.72 | 0.003162 | LPE(18:1) | -1.36 | 0.000701 |
| TG(O-18:1_16:0_18:1) | -1.65 | 0.001677 | LPE(18:2) | -1.68 | 0.000701 |
| TG(18:1_20:4_22:6) | 1.74 | 0.006201 | PC(18:0_22:6) | 1.27 | 0.004289 |
| TG(18:1_21:0_16:0) | -1.05 | 0.006201 | PE(16:0_18:2) | 1.11 | 0.000449 |
| TG(18:1_22:0_18:1) | -1.97 | 0.006618 | PE(16:0_20:4) | 1.08 | 0.000832 |
| TG(18:1_22:0_22:0) | -1.8 | 0.002948 | PE(16:0_20:5) | 1.44 | 0.005871 |
| TG(O-18:2_18:1_18:1) | -1.48 | 0.005676 | PE(16:0_22:6) | 1.27 | 0.000832 |
| TG(18:2_20:4_22:6) | 1.7 | 0.004729 | PE(18:0_20:5) | 1.06 | 0.004658 |
| TG(18:2_22:6_22:6) | 2.06 | 0.004839 | PE(18:0_22:6) | 1.56 | 0.000608 |
| TG(O-20:0_18:1_18:2) | -1.19 | 0.008578 | PE(18:1_22:6) | 1.2 | 0.000455 |

|  |  |  |
| --- | --- | --- |
| TG(24:1_18:2_18:1) | -1.87 | 0.008578 |

Neutral lipids

In PbA-MV, all PE species were higher than control except PE(O-16:1_18:2) and PE(O-18:1_18:2) (Table I). Compared to MV from control mice, Py-MV had only three PE lipid species that were significantly increased (Table II), in contrast to the twelve that were increased in PbA-MV. When comparing MV from PbA-infected to those from Py-infected mice, we found 7 PE lipid species increased and two decreased (Table III). No LPE species were significantly modulated in Py-MV (Table II). In contrast, LPE(18:1) and LPE(18:2) were reduced in both PbA-MV vs control (Table I) and vs Py-MV (Table III).

A striking number of identified LPC lipid species were significantly reduced in PbA-MV vs control-MV and vs Py-MV, but there was no difference in the Py-MV vs control-MV. Twelve PC species were significantly modulated in PbA-MV vs control-MV (Table I) and Py-MV showed only two PC species in lower amounts than in controls (Table II), despite no significant difference in total PC amounts (Fig. 2). Interestingly, PC(18:0_22:6) was higher in PbA-MV compared to both control-MV and Py-MV.

PS(18:0_22:6) lipid amounts were higher in both PbA- and Py-MV vs controls (Table I & II), while two additional PS lipid species were significantly modulated in Py-MV vs control (Table II). With regard to Hex1Cer lipid species, both Hex1Cer(d18:1_16:0) and Hex1Cer(d42:1) species were higher in PbA- and in Py-MV than in controls (Table I & II). However, Hex1Cer(d41:1) was significantly lower than control in PbA-MV only (Table I). SM(d36:0) was significantly increased, and PI(O-34:3) was significantly decreased compared to control only in PbA-MV (Table I). In the DG class, six species were found to be significantly higher in PbA-MV than in controls (Table I), and three were significantly higher in PbA-MV than in Py-MV (Table III). No DG lipids were significantly modulated between Py and control (Table II). No changes were observed in ChE and Cer lipids in our study.

Finally, we detected TG lipids in the MV preparations, and we comprehensively characterised the fatty acyl chain composition of the lipid species. Despite the lack of difference between the three groups at the TG class level (Fig. 2), numerous TG species were differentially expressed (Table I and Fig. 4). Interestingly, the five TG species containing docosahexaenoic acid (DHA FA 22:6) in their composition were the most strikingly different: all of them were higher, while all other identified TGs were lower, in PbA-MV than in both Py- and control-MV, as highlighted in yellow boxes (Fig. 4). Remarkably, there was no difference in any identified TG species between Py-MV and controls.

**Fig. 4.**
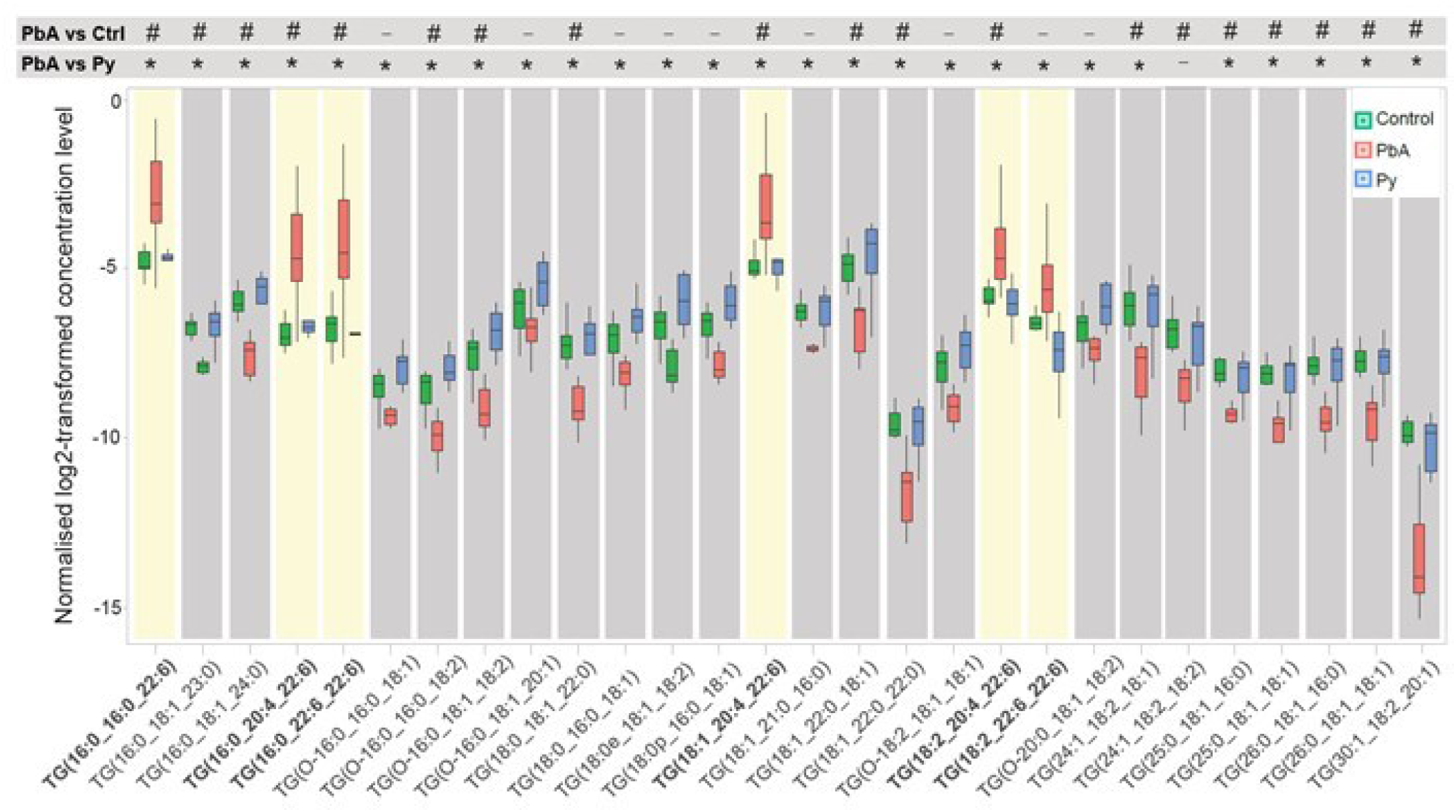
Docosahexanoeic acid (DHA 22:6) containing TG lipids are higher in plasma MVs during cerebral malaria (CM). Comparison of levels of characterised TG species from plasma MVs between PbA (red) vs control (green) and Py (blue). Yellow boxes highlight DHA 22:6 containing TG lipids that are exclusively elevated in PbA vs control and Py. To display the statistical significance of differences, symbols (-) represents no statistical difference, (#) for PbA vs control, and (*) for PbA vs Py based on adjusted p-value <0.01. There are no differences in TGs between Py and control.

The correlation heatmap (Fig. 5) shows pairwise correlations among 121 lipids retained after removing invariant lipids (IQR ≤ 1) across all the samples. Changes are either in concordance (positive correlation shown in blue) or inverse concordance (negative correlation shown in red). Hierarchical clustering of the complete lipid-lipid correlation matrix describing 14 520 unique pairs of lipids (excluding self) revealed eight distinct clusters of positively correlated lipids (Pearson correlation > 0.7), organised along the diagonal of the matrix. Clusters 1 and 3 are highly expressed in PbA vs control or vs Py and essentially are composed of TGs and DGs. In contrast, cluster 2 is composed of lipids that are increased in PbA- and Py-MV vs control, but not between PbA- and Py-MV (Fig. 5A). All other clusters, excluding cluster 8, are lowly expressed in PbA vs control/ Py. Interestingly, cluster 8 is increased, is composed mostly of Hex1Cer, PE and PS and has a similar change in both PbA- and Py- vs control-MV. Cluster 3 shows TG and DG lipids that are composed of 22:6 in their structure and have positive fold change in PbA- vs control-MV, which is potentially related to CM pathogenesis. Cluster 3 shows negative correlation with cluster 4, which is mainly composed of phospholipids, which are lowly expressed in PbA- vs control- and Py-MV (Fig. 5A). Clusters 6 and 7 are the largest, with cluster 6 composed mainly of TGs and cluster 7 composed mostly of PC and LPC, both with negative fold change in PbA- vs control-MV and vs Py-MV and could play a role in CM phenotype progression that is opposite to that of TG containing 22:6 in cluster 3.

**Fig. 5.**
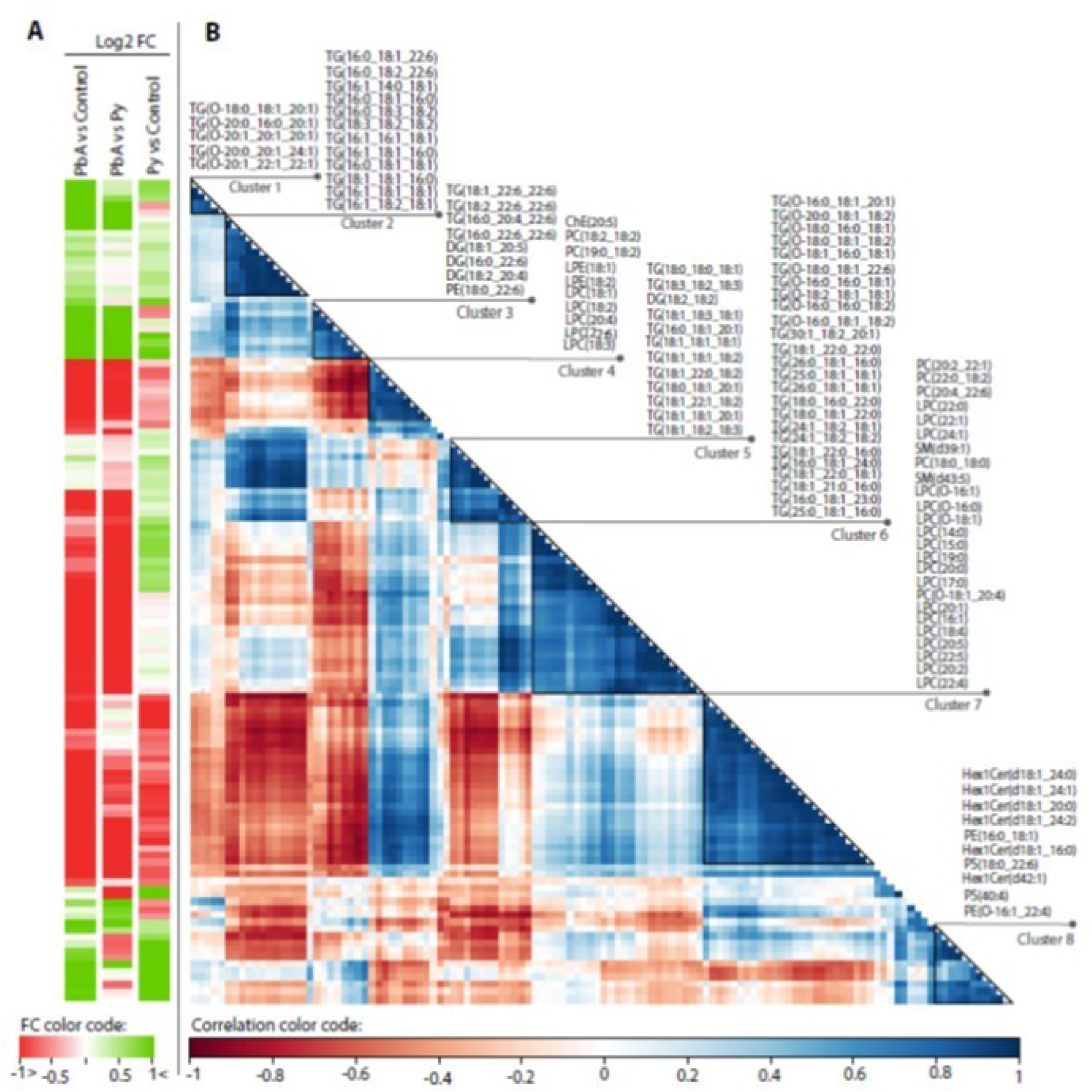
Correlation plot of the 121 retained lipids across all the samples. (A) fold changes (FC) of lipid amounts in the three categories of comparisons. Green denotes an increase and red a decrease. (B) Hierarchical clustering of the lipid-lipid correlation matrix. Rows and columns correspond to the 121 retained lipid species. Dark blue triangles indicate clusters (1-8) of strongly positively correlated lipids. Cluster numbers and corresponding lipid names are shown on the right.

## Discussion

In this paper, we identified unique profiles of MV lipid species/classes in relation to CM. In particular, total LPC levels were significantly lower in PbA- and Py-MV compared to uninfected mice and in PbA- compared to Py-MVs. Increased concentrations of circulating LPC is associated with inflammatory disorders^37^, and more recently has been found in increased amounts in platelet MV in myocardial infarction and atherosclerosis^38^. Total LPE levels were also lower in PbA compared to uninfected mice and Py-infected mice. Based on current knowledge, the reduction in both LPE and LPC in CM MVs suggests a lack of inflammation. However, these differences observed in plasma MV between PbA and Py infection are consistent with the strong immunopathological response underpinning CM, as opposed to non-CM, as detailed in several works^4,21,39-41^ and warrant additional investigation. Activation of phospholipase A 2 (PLA2) is required to cleave the fatty acid on sn2 position in phospholipids; this most commonly releases arachidonic acid, which is then converted to inflammatory eicosanoids. The other product is lysophospholipids such as LPC and LPE. Pappa *et al*. showed a positive correlation between PLA2 activity and neural inflammation in children with CM^42^. Investigating PLA2 levels and activity in CM MVs is of interest to elucidate if it plays a role in modulating the levels of MV lysophospholipids and consequently the development of CM.

PE levels were significantly higher in MV from PbA mice compared to those from uninfected control and Py. On the other hand, total PS levels were significantly higher in both PbA and Py compared to uninfected control, suggesting a role for PS in malarial infection but not CM pathogenesis specifically.

DHA 22:6 have a known role in inflammation as they belong to the ω-3 pathway^43^ and thereby are anti-inflammatory. In our study, 13 out of 16 DHA (22:6) containing lipids were found to be increased, while they were decreased in 2 out of 16 lipids in PbA-MV vs control-MV, suggesting an anti-inflammatory potential in MV circulating at the time of CM. Conversely, in MV from mice with non-CM, 2 of these were up-regulated and 2 were down-regulated, when compared to MV from control mice. Although DHA plays a protective role in inflammation, deleterious effects have also been reported, where an excess of ω-3 membrane lipids can increase the susceptibility to infection^44^.

It has been suggested that high levels of TG lipids in MV preparations are considered as an indication of contamination from other vesicle types present in plasma^45^. Surprisingly, despite the total level of detected TGs in our preparations being comparable between the three types of MV populations, TG lipids containing DHA 22:6 were exclusively increased in MV preparations from PbA infected mice, indicating the potential utility of these lipids to predict CM development in mice.

Given that MVs are taken up by macrophages, it is tempting to speculate that the changed lipidome of MVs during CM plays an anti-inflammatory role or is a mechanism used by the parasite to modulate the host immune response^41^. More specifically, it is possible to hypothesise that MVs utilise TGs to supply DHA 22:6 to recipient cells that incorporate it into membrane phospholipids. Whether the role of DHA is deleterious (by increasing membrane fluidity or affect the MV ability to fuse with target cells) or protective (by anti-inflammatory properties) remains to be elucidated.

Correlation maps showed clusters of lipid species which changed together in a related fashion; for instance, the changes in TG and DG correlated, which is not surprising since they are biologicaly linked sharing similar biosynthetic pathways. The possibility that TG is being used as a storage site for fatty acids that can later be hydrolysed and employed for synthesis of additional lipids remains of interest for future investigation.

In summary, these results suggest that experimental CM is characterised by specific changes in lipid composition of circulating MV. More specifically, the key differences between CM and non-CM MVs characterised by the reduction in LPC and LPE and a specific increase in TGs conatining DHA containing TGs. Microvesicles carry a large array of active molecules including lipid mediators, phospholipases, proteins and RNA that can be used to modulate the phenotype of recipient cells^46^. Future studies investigating differences in lipid packaging and phospholipase activity in microvesicles from CM vs NCM, may shed some light on the role these lipids play in malaria complications.

## Acknowledgements

This work was funded by the National Health & Medical Research Council of Australia (grant #1099920 to GEG and NHH). BCAL Dx provided financial support for mass spectrometry analysis and AB salary. We thank Dr David Peake and MKI for providing us LipidSearch support. We thank Kerry Heffernan for technical support and review of the manuscript.

## Author Contributions

GEG, AB and NHH designed the study. AJ, AC and EHB performed the in vivo and MV preparation experiments. AB conducted lipidomics experiments including LCMS method development and analysis. AB carried out the data analysis. AB, FV and GEG carried out the statistical analysis. AB and FV carried out data visualization. AB and GEG wrote the manuscript. TM critically reviewed the manuscript, provided support on presenting the data and assisted with writing the manuscript. AC and MM provided technical and software support, LCMS method assistance, instrument support critical for conducting this work. All authors revised the manuscript.

