## Supplemental Figure 1 and 2 for "Lipidome profiles of plasma microvesicles differ in experimental cerebral malaria, compared to malaria without neurological complications"

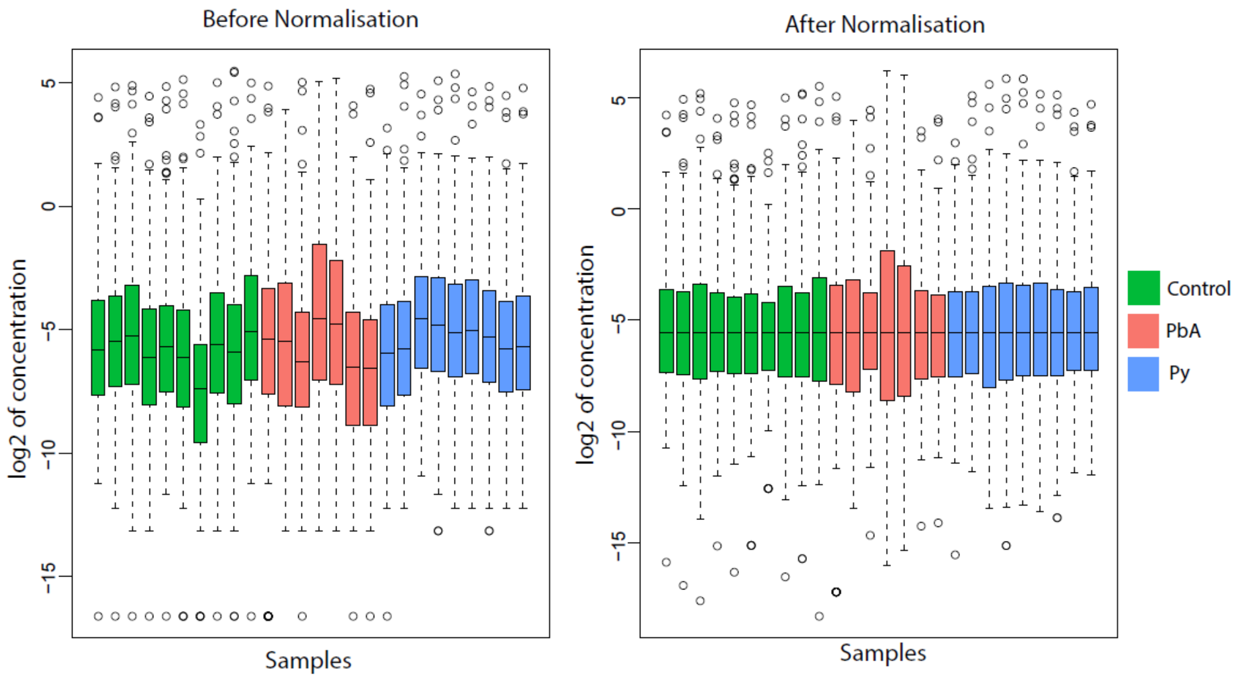


Supplementary Fig.1. **Boxplots of samples before and after median normalisation**. Each graph represents the minimum, maximum, median, first quartile and third quartile in the data set. Circles are outliers. An outlier is defined as a data point that is located outside 1.5 times the interquartile range above the upper quartile and bellow the lower quartile

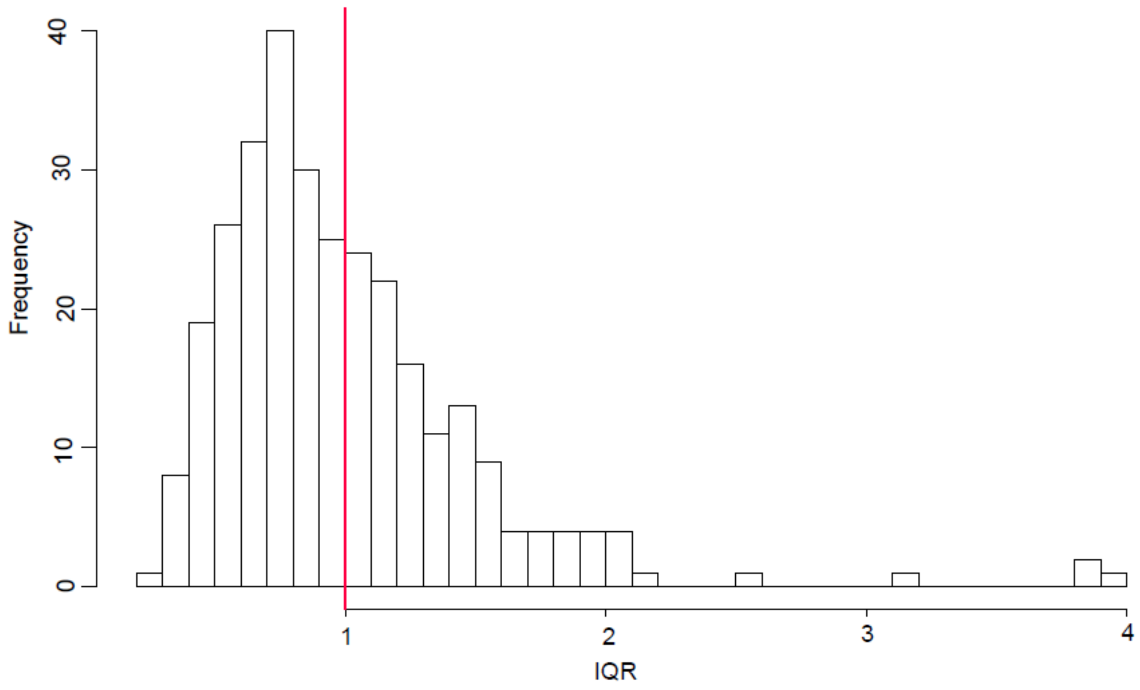


Supplementary Fig. 2. **Histogram of Interquartile range (IQR) representing variability of lipids across samples.** Lipids with IQR ≤ 1 (red line) were considered as invariant and removed before further analysis
